# Mechanistic model: Analysis of diffusion dynamics of Lugol’s chromoendoscopy in Basal layer-type squamous cell carcinoma in situ via Euler’s method and Law of mass action

**DOI:** 10.1101/2021.05.13.444011

**Authors:** Fei Xiong

**Affiliations:** Sichuan Provincial People’s Hospital, University of electronic science and technology of China, Chengdu, Sichuan, PR China

## Abstract

Basal layer-type squamous cell carcinoma in situ(BLSCC) is an aggressive malignancy tumor. In the past, it usually went ignored because its aggressive biological behavior is different from typical esophageal squamous cancer(SCC). For our mechanistic model, we analyzed diffusion dynamics of iodine stain in BLSCC with the help of Euler’s method. We used the law of mass action description of the forming process of the iodine-glycogen Complex. Based on the above analysis, our mechanistic model reveals that the efficacy of Lugol’s chromoendoscopy is determined by the thickness of the squamous epithelial cell layer, which contains glycogen granules, and the convection of the interpupillary capillary loop(IPCL). A further inference can be made: the unique histological structure of BLSCC suggests Lugol’s chromoendoscopy does not detect some components of it because these components neither break enough squamous epithelial cells which contain glycogen granules nor contain enough number of IPCL in a specific time frame. Besides, if our mechanistic model starting point is BLSCC and typical SCC share the same evolution path, we found both their cytoplasm tends to solid phase from the iodine diffusion dynamics analysis in “pink color sign.”

## Introduction

With the evolution of endoscopic technique, there has been a change in the stage distribution of esophageal cancer. The incidence rate of superficial esophageal cancer is increasing over the last two decades(Kanamoto, Yamaguchi et al. 2000, Wang, Jiao et al. 2004). According to recent studies, The risk of nodal metastases of early esophageal cancers (invading no deeper than SM1) is slower than beyond the depth of invasion SM1(Manner, Pech et al. 2013, Cho, Choi et al. 2014). Early esophageal cancer has high cure rates because of low lymph node metastasis rate(Funakawa, Uto et al. 2015), so early esophageal cancer screening becomes more significant.

The distinction of early esophageal neoplastic changes from normal esophageal mucosa can be a difficult task for conventional endoscopy(Hashimoto, Iriya et al. 2005). Chromoendoscopy with iodine staining has a higher sensitivity than conventional endoscopy(Dawsey, Fleischer et al. 1998, Freitag, Barros et al. 1999), so it becomes a good choice for detecting early neoplastic changes. It is well known that Iodine’s esophageal mucosa absorption mechanism is passive diffusion(Ishihara, Kanzaki et al. 2013). Common squamous epithelium stains dark brown (Figure A) because glycogen with iodine union inside cells can produce this colour. These molecules are held together by electrostatic forces(Moulay 2013), so it is a reversible process. Yellower color, even pink color, is usually associated with Neoplastic cells because there is no glycogen in these tumor cells(Ishihara, Kanzaki et al. 2013) (Figure B). However, the new definition of High-grade dysplasia from WHO(Board 2019) suggests that Chromendoscopy with iodine staining may be less effective than the past understandings of superficial esophageal cancer. The newest version features slight modification to the definition: High grade involves more than half of the epithelium and severe cytological atypia regardless of the extent of epithelial involvement. The latter implies that high-grade dysplasia is not more than half of the epithelium(Figure D) and brings many difficulties in its diagnose and treatment(Sarbia, Wolfer et al. 2015). Japanese pathologists called it Basal layer-type squamous cell carcinoma in situ(Zhuang, Chen et al. 2019) (BLSCC).compare with the conventional carcinogenesis way of typical SCC, BLSCC infiltrates directly to lamina propria and do not involve in the upper half of the epithelium. Chromoendoscopy with iodine staining may be more reliable than narrow-band imaging in detecting BLSCC because the latter is harder to detect irregularly shaped multiform lesions and absence of homogenization(Ishihara, Takeuchi et al. 2010, Goda, Dobashi et al. 2015, Zhuang, Chen et al. 2019). However, Xiaoduan Zhuang et al. showed. Basal layer-type squamous cell carcinoma in situ had a higher recurrence rate(Zhuang, Chen et al. 2019), so BLSCC’s recurrence rate implied some limitations of Lugol’s Chromendoscopy for detecting a demarcation line between normal mucosa and cancerous lesion. More interesting, For BLSCC, scattered stained spots in Iodine unstained areas(Zhuang, Chen et al. 2019) suggested there exist two color patterns of Lugol’s Chromendoscopy: yellow and brown-dark. The latter color is usually considered a normal mucous membrane in the past(Ishihara, Kanzaki et al. 2013). In short, there are two problems with Lugol’s Iodine: 1.Do Lugol’s Chromendoscopy have the ability to detect a demarcation line between normal mucosa and BLSCC? 2. Why does exist such a difference in Lugol’s Iodine staining patterns in BLSCC? difficult diagnosis(Zhuang, Chen et al. 2019) may lead to statistical models based on data from endoscopic technique and pathology, even next-generation sequencing, do not fully answer the first problem; further, the quantification of data’s randomness(Agresti and Franklin 2018) is not enough to explain the second problem and the mechanism hidden behind. Understanding the mechanism hidden behind it can help develop more effective diagnostic and therapeutic strategies; thus, our problems need to be explained differently: a mechanistic model based on the histological structure.

**Figure A.**
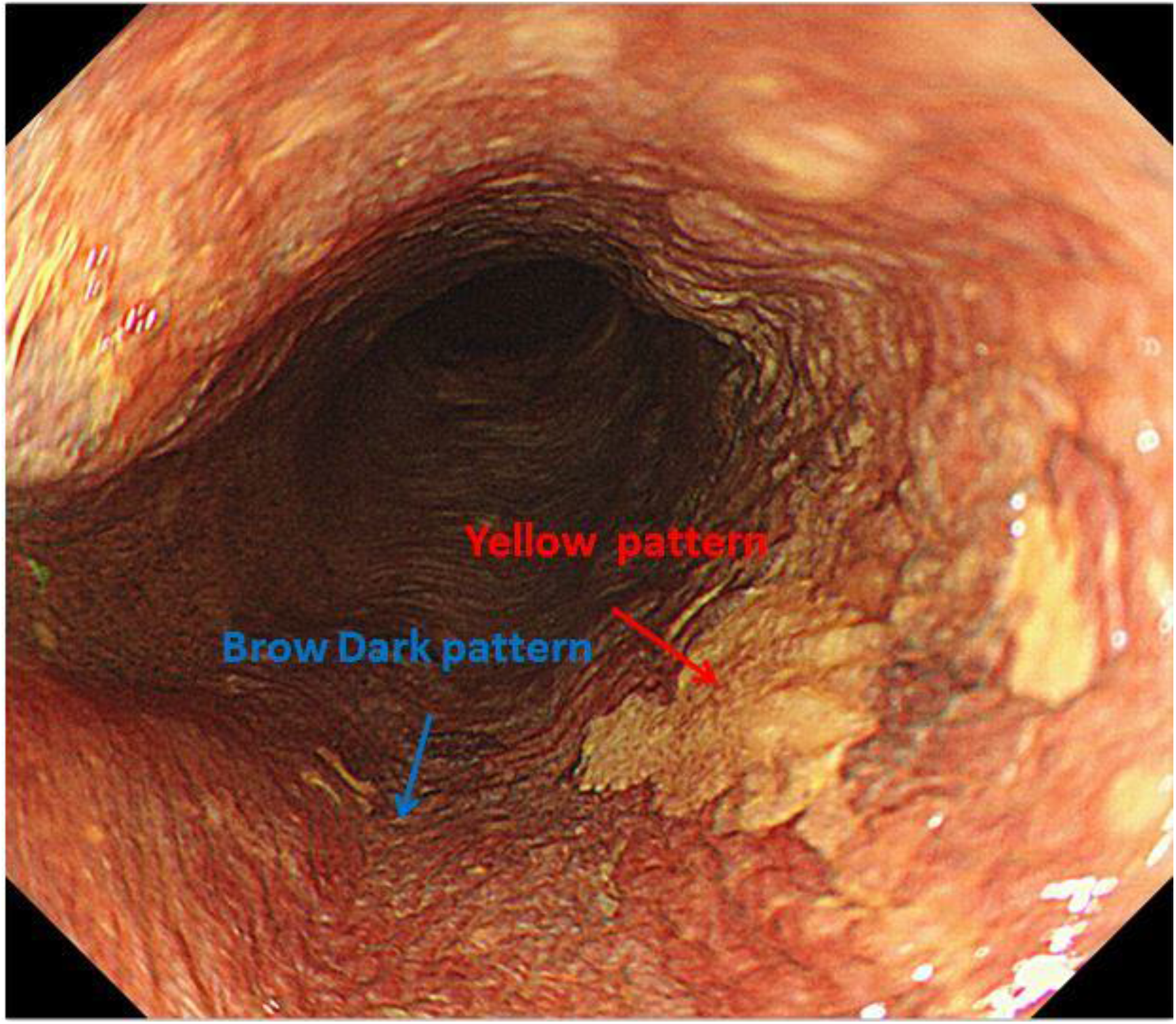
Lugol’s chromoendoscopy: 1.Brow dark pattern is normal squamous epithelium; 2. Yellow pattern is usually associate with dysplasia or early malignancy.

**Figure B.**
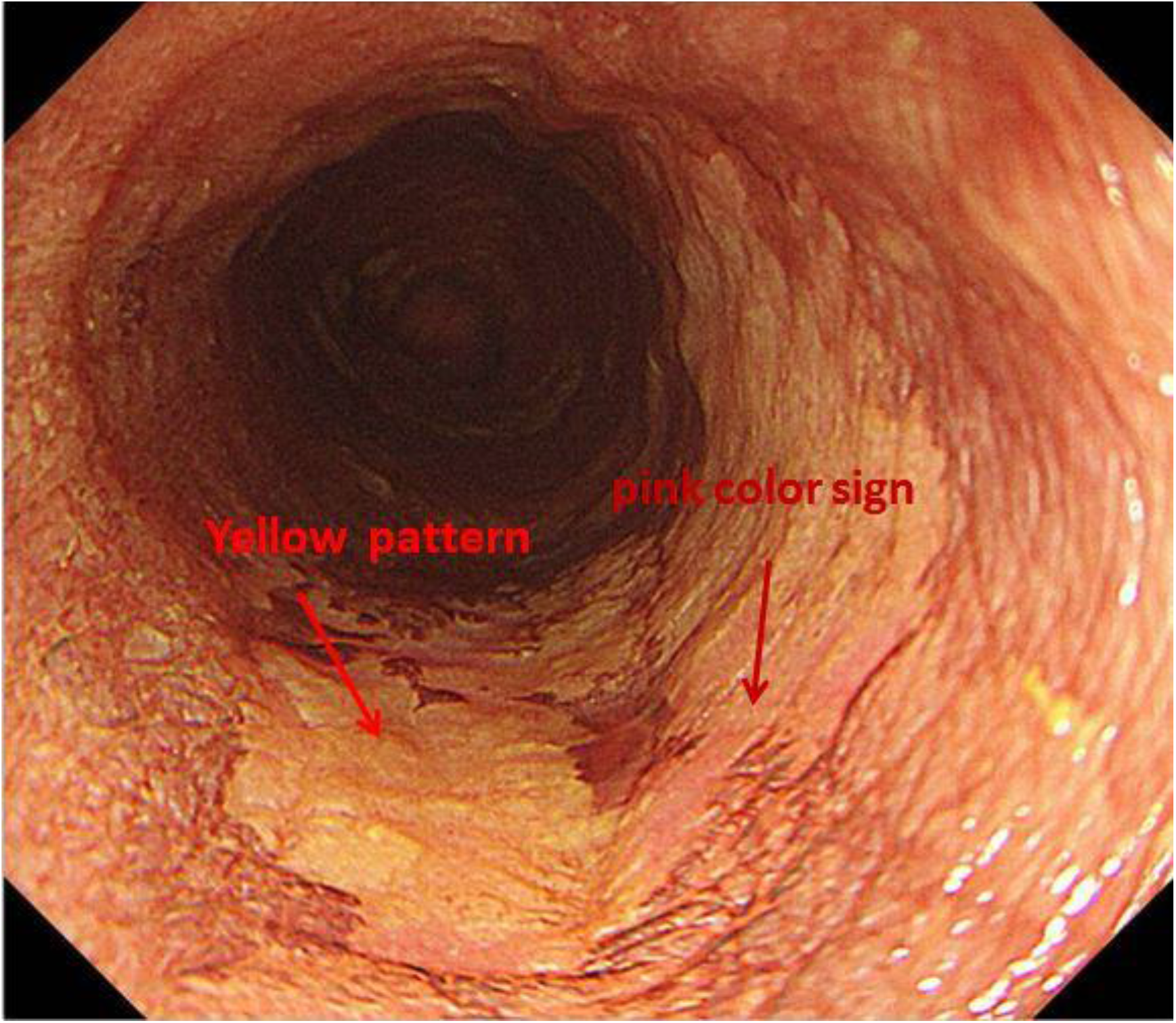
“pink color sign” means high grade dysplasia involves the whole epithelium layer, Yellow pattern dysplasia don’t involve the whole epithelium layer.

**Figure C.**
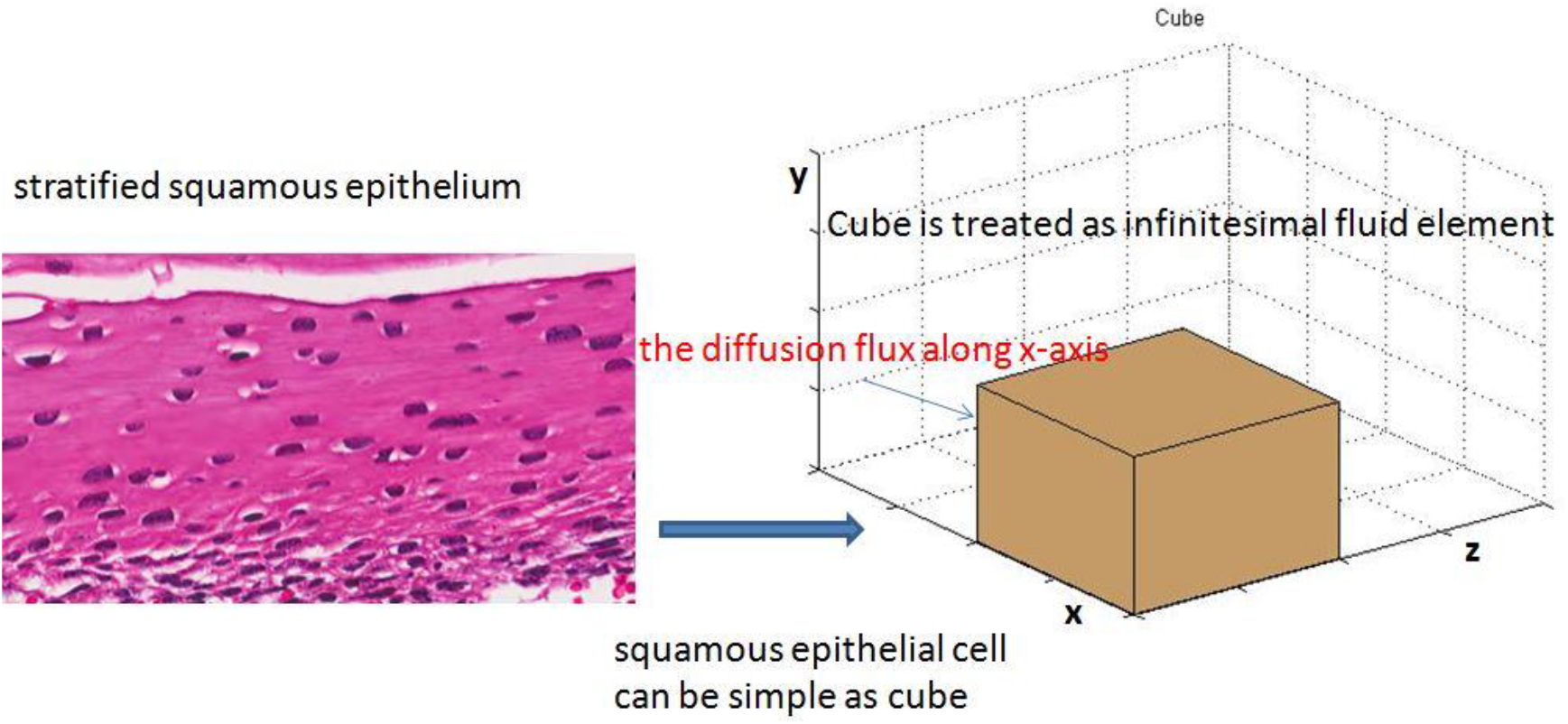
For our model, the nonkeratinizing, stratified squamous epithelium can be as simple as multilayer cubes and infinitesimal fluid elements; it is clear that the calculation of diffusion direction of Iodine is simplified.

**Figure D.**
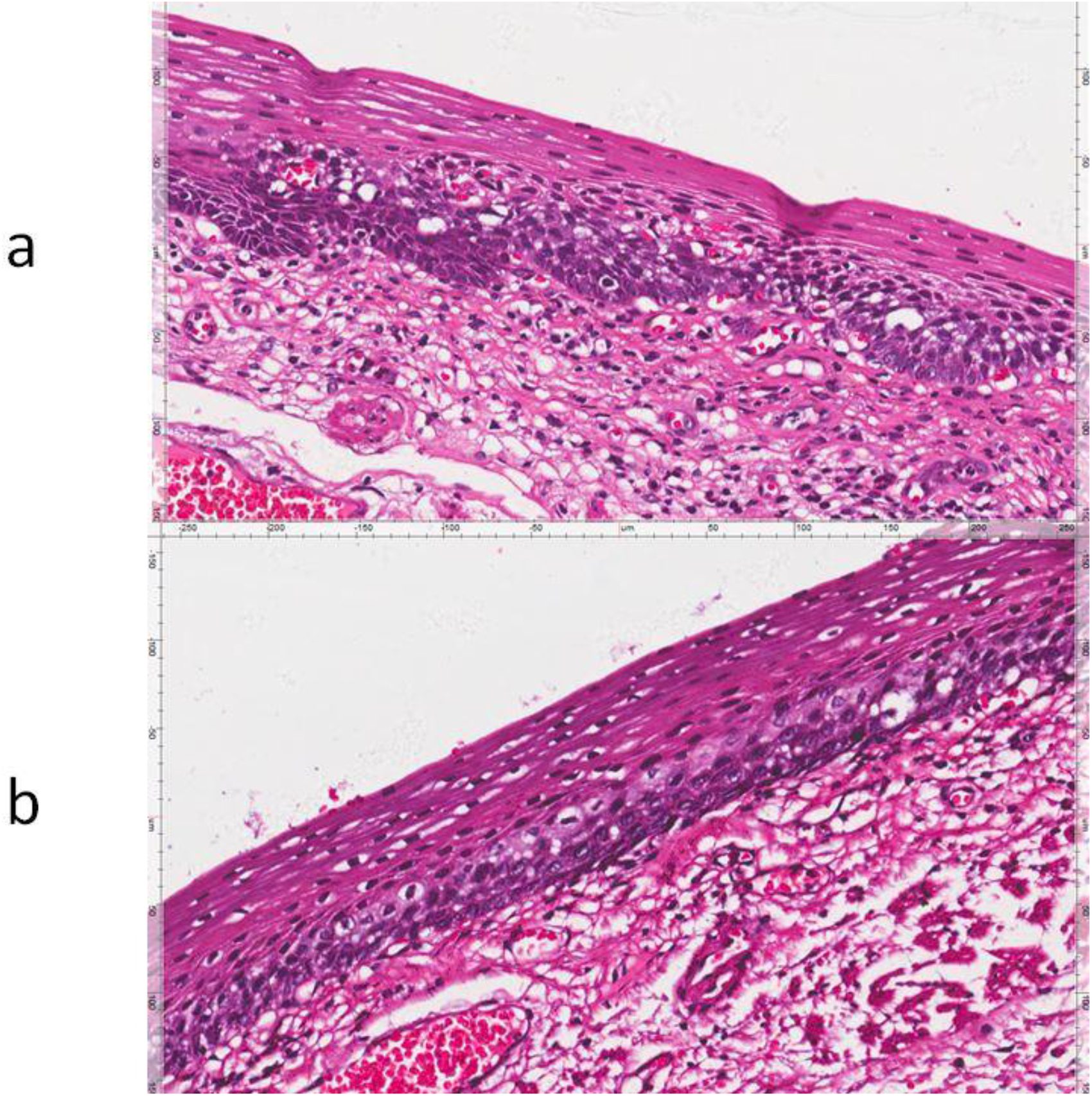
FigureD(a) and (b) shows BLSCC’s unique histological structure. It is similar to the definition of high-grade dysplasia from WHO(2019): severe cytological atypia regardless of the extent of epithelial involvement. Please note the difference between (a) and (b). Due to tumor-associated angiogenesis of BLSCC, the integrated effects of the maximum distance of nutrient diffusion and the degree of cytological atypia may be determined. (Note: The picture does not show the actual thickness of epithelium because the specimen of endoscopic submucosal dissection tends to shrink by dehydrated)

## Model

Although the squamous epithelium is composed mainly of three cell layers(Mills 2012), each cell could be seen as a flat cell from morphological characteristics. We abstract the flat cell into a cube because the diffusion direction of Iodine is restricted in x,y, z-axis in 3D dimensional space(Figure C). It is clear that the cube reduces the computational complexity of diffusion direction, so the nonkeratinizing, stratified squamous epithelium can be as simple as multilayer cubes(Figure C). We removed the cell nucleus and other organelles for the same principle because there is no relationship between them and iodine diffusion. As mentioned above, Basal layer-type squamous cell carcinoma in situ is High-grade dysplasia, restricted to the basal cell layer(Zhuang, Chen et al. 2019) (Figure D). In our model, we use a black cube for BLSCC(Figure E). Come back to our problems; we need to solve the demarcation line problem, so the key of our model is that tumor and normal tissues are joined in step base on the histology of BLSCC(Figure E). The molecule iodine diffuses to the glycogen interface forms an Iodine- glycogen complex(Ishihara, Kanzaki et al. 2013, Moulay 2013). To simplify our schematic diagram, we choose to ignore it and deduce the process by Law of mass action(Ferner and Aronson 2016)

**Figure E.**
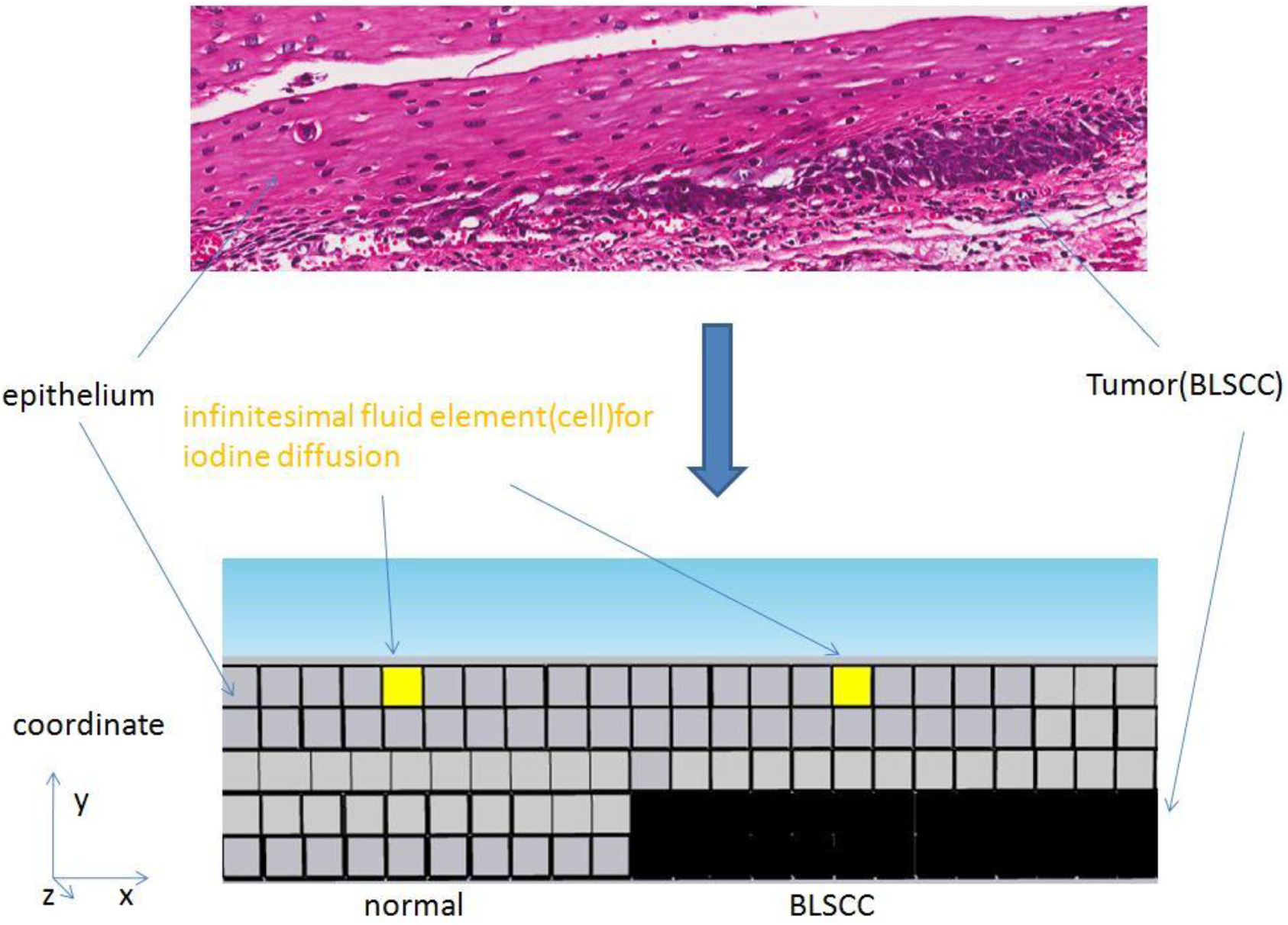
Tumor and normal tissues are joined in step base on the histology of BLSCC. BLSCC are black cubes; we analyze the iodine diffusion between BLSCC and normal epithelium via the Euler method through the geometry structure.

## Analysis and Result

Recent researches have shown that liquid-liquid phase separation does exist in the cytoplasm(Alberti, Gladfelter et al. 2019), and Macromolecular crowd is a common phenomenon inside the cell(Mourão, Hakim et al. 2014), it is evident that the viscous force exists in the natural fluid inside the cell. However, at the tissue level, the effect of it is almost imperceptible on Lugol’s Chromendoscopy, more critical, the inputs and outputs of Iodine are both passive diffusion, which suggests the free diffusion dynamic of Iodine has continuity characteristics, so we ignore the viscosity and make an approximation which describes “dry water”(Feynman, Leighton et al. 1963) rather than natural state inside the cell. According to Beer-Lambert law(Mayerhöfer, Pahlow et al. 2020), we have

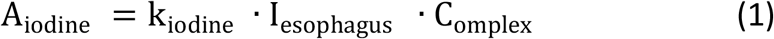

where A_iodine_ is the absorbance of normal esophageal mucosa after iodine staining;k_iodine_ is attenuation is attenuation coefficient of normal esophageal mucosa after iodine staining; C_omplex_ is the density of Iodine- glycogen complex; l_esophagus_ is the length of esophageal epithelium which contain glycogen granules along the y-axis(note: let us first assume that C_omplex_ is low concentration in our model).

A single cell’s model (cubic), suppose that the C_complex_term mainly determine the color pattern. The Charge-transfer complex(Moulay 2013) means there is a chemical equilibrium between Iodine and glycogen:

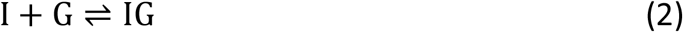

where I is Iodine; G is glycogen; IG is Iodine- glycogen complex. (note: although Lugol iodine is a solution of iodine ions with Iodine, the organic reaction occurs only in the presence of a single iodine molecule. To simplify our complex reaction, we take Lugol iodine as a solution of Iodine in our model)

For a single cell (cubic), It is clear that G is constant and I is variable. Eq(1), k_iodine_ and l_esophagus_ are fixed in a single cubic; logically, A_iodine_ is determined by the density of Iodine. The density of Iodine represents the state of every spatial point in iodine diffusion at some point, so Euler’s method(Anderson and Wendt 1995) is very suitable to use in our model.

It is well known that Iodine across the cell membrane by passive transport(Ishihara, Kanzaki et al. 2013).the movement of Iodine usually across cytomembrane via facilitated diffusion mechanism(E Nunez-Anita, Cajero-Juarez et al. 2011) it is generally recognized that the thickness of the cell membrane is approximately 3nm(Van Meer, Voelker et al. 2008), The volume of it takes fractional percentage point of the volume of the cell. Let us assume that number of iodine molecules is less than the transport protein on the cell membrane and Iodine continually diffuse into an infinitesimal fluid element. We can consider the transport cycle as an instantaneous state relative to the macro world to ignore the cytomembrane in the diffusion process of Iodine.

According to the oxygen diffusion mathematical model(Endeward 2012), the rate of iodine concentration is appropriate for Fick’s second law in the infinitesimal fluid element during the process of facilitated diffusion. Now we think of a single cell (cube) as an infinitesimal fluid element (iodines) (Figure C). For the x-axis, suppose that the concentration gradient is 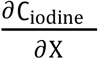 at cubic’s top edge; so we have:

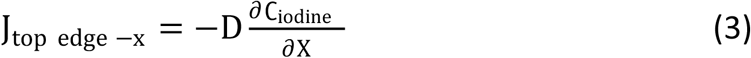

where J_top edge −x_ is the diffusion flux at the cubic top edge along the x-axis; D is diffusivity; **C**_Iodine_ is the concentration of Iodine at cubic top edge; X is position, the dimension of x-axis; 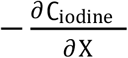 is the concentration gradient of Iodine for one dimension diffusion at cubic top edge.

Then, let us assume that the concentration variation rate of our cubic is continuous, so we use the Taylor polynomial(Thomas, Weir et al. 2008) to approach the real function of the concentration gradient at the cubic bottom edge:

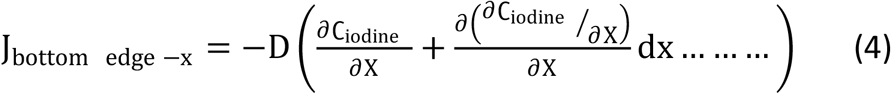

where J_bottom edge −x_ is the diffusion flux at cubic’s bottom edge along the x-axis; 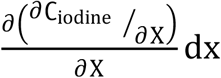 is the second term of Taylor’s expansion and equal to Fick’s second law.

For tissue level, the width of the cube is infinitesimal, so we have

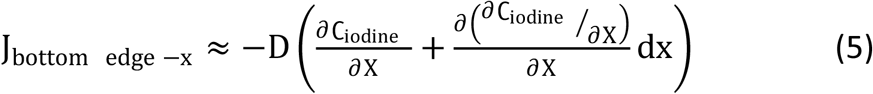

Then, according to mass conservation(Feynman, Leighton et al. 1963), the resultant diffusion flux at cube along the x-axis is

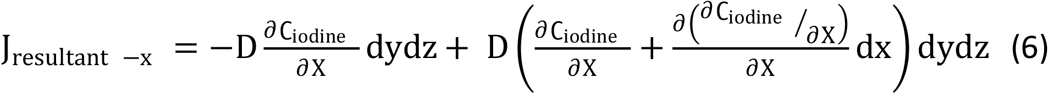

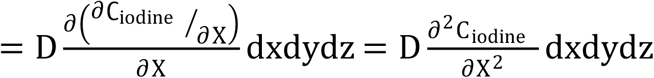

where J_resultant −x_ is the resultant diffusion flux at cubic along the x-axis.

Similarly, the resultant diffusion flux at cubic along y-axis and z-axis are:

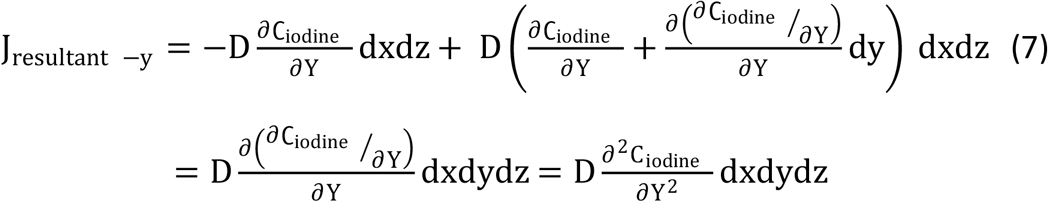

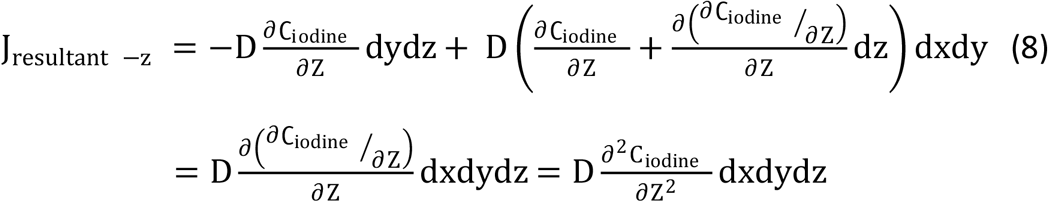

where J_resultant −y_ and J_resultant −z_ are the resultant diffusion flux at cubic along y-axis and z-axis, respectively.

Summing Eq(6),Eq(7) and Eq(8),we get

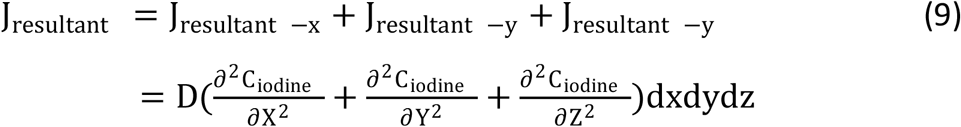

where J_resultant_ is the resultant diffusion flux at cubic in the ideal state. Eq(9) shows a total disregard for an organic reaction between Iodine and glycogen(Moulay 2013), so we write Eq(9) as:

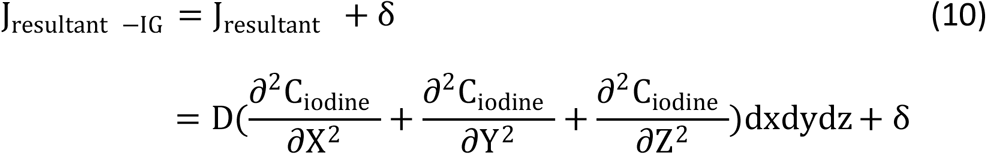

where δ impacts the organic reaction between Iodine and glycogen resultant diffusion flux at cubic. J_resultant −IG_ is the resultant diffusion flux at cubic involves the organic reaction between Iodine and glycogen.

Now consider how the organic reaction takes effect on J_resultant −IG_. According to Eq(2), there are only two states of Iodine in our model: iodine molecule and Iodine- glycogen complex. For the total concentration of Iodine in our model:

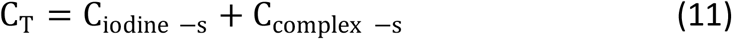

Where C_T_ is the total concentration of iodine molecule and Iodine glycogen complex at single cube. C_iodine −s_ is the concentration of iodine molecules at a single cube. C_complex −s_ is the concentration of Iodine- glycogen complex at a single cube.

According to the Brownian movement theory(Feynman, Leighton et al. 1963), the iodine-glycogen Complex comes from a random collision between Iodine and glycogen. Law of mass action(Ferner and Aronson 2016) can be used to describe the process:

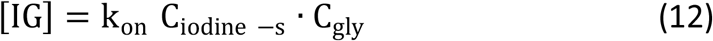

where [IG] is the formation rate of Iodine- glycogen complex;C_gly_ is the concentration of unconjugated glycogen at a single cubic; k_on_ is the collision rate between Iodine and glycogen(note: it means how many collision events are happening per single Iodine molecular per second, when the total concentration of glycogen is constant).

Similarly, k_off_ represents the dissociate rate of Iodine- glycogen complex. Normally, the formation heat of the Complex is −40kj/mol(Moulay 2013), so k_off_ is a positive constant. Base on the processes of dissociating and collision, we have:

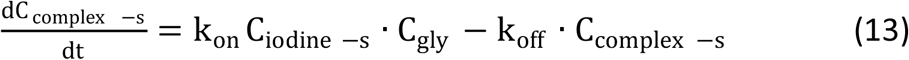

Eq(13) is finally close to the static equilibrium. For chemical equilibrium, we have:

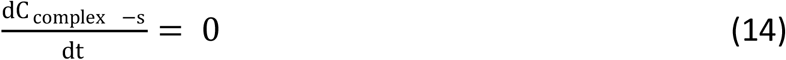

Using Eq(13) with Eq(14), we get:

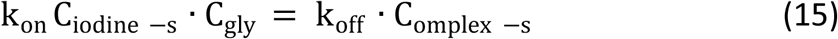

The definition of k_on_ tells us k_on_ is not a constant because C_gly_ is variable during the diffusion process of Iodine. From the point of view of statistics, there is positive relativity between two variables base on collision probability.

If we combine Eq(15) with Eq(11), we get:

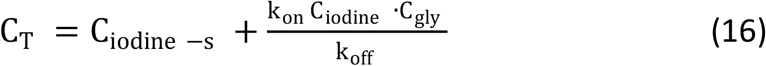

So we can say that for C_iodine_:

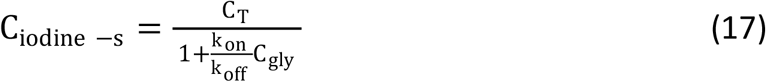

Before we discuss the dynamics of Eq(18), we need to know: if C_T_ approach +∞, does the organic reaction between Iodine and glycogen always keep chemical equilibrium?

We can explain the problem above with the help of proof of contraposition(Rosen 2011). If C_T_ approach +∞, suppose that the chemical equilibrium is broken. 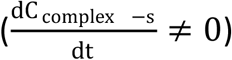The infinite value of C_omplex_ suggest all glycogen react with Iodine, so we get:

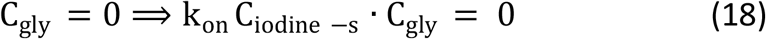

So we write Eq(13) as:

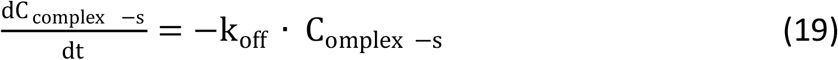

By integrating both sides of Eq(19):

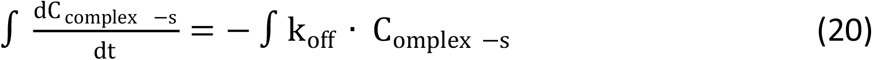

So we obtain:

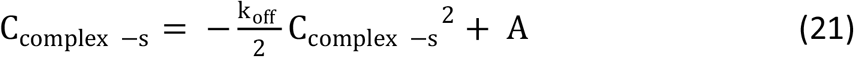

where A is Constant of integration.

Let all iodine molecules at the top edge of cubic in our model, we have:

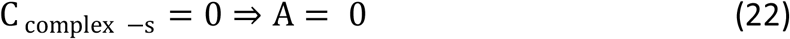

So that Eq(21) becomes:

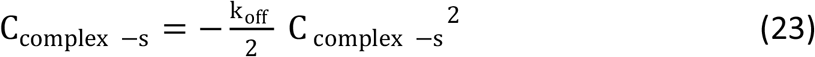

Substituting the equation in Eq(11), we find that:

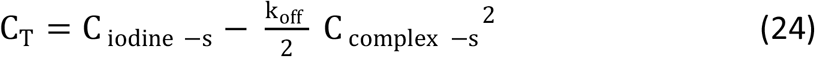

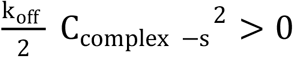, Eq(24) means:

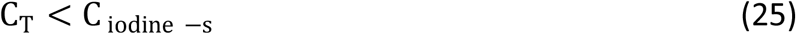

However, according to Eq(11), we have:

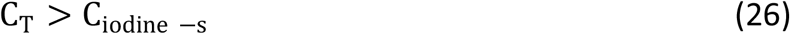

These two equations are mutually incompatible. Indeed, the hypothesis “if C_T_ approach +∞, the organic reaction between iodine and glycogen do not keep chemical equilibrium” is false. So “the organic reaction always keeps chemical equilibrium” is automatically true.

Let us assume that the total concentration of glycogen is constant, so we get:

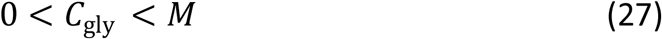

where M is the constant(the total concentration of glycogen before the chemical reaction occurs).

Eq(2) tell us:

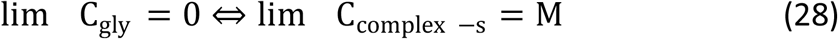

Let us assume that C_omplex −s_ approach M after the color of the esophageal mucosa turns into brown-dark. Besides, According to the definition of k_on_, k_on_ is determined by the value of C_gly_, which means both of them to have the same monotonicity, so we get:

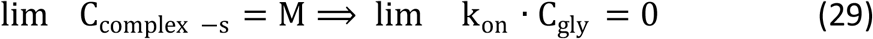

Therefore, we can rewrite Eq(17) as:

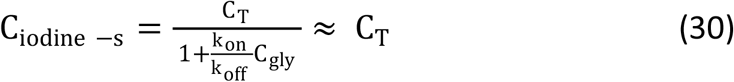

According to the definition of resultant diffusion flux, we get:

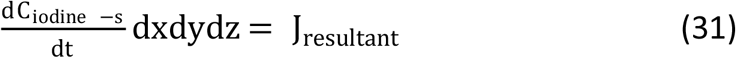

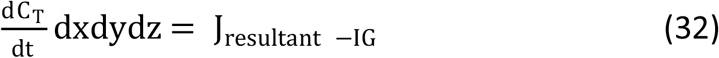

Therefore, differentiate both side of Eq(30) and multiply by dxdydz: J_resultant −IG_ ≈ J_resultant_

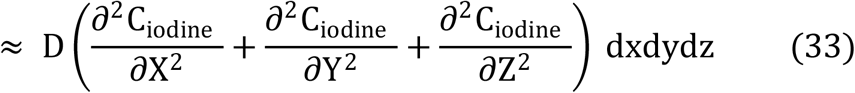

The equation says that for C_omplex −s_ approach M the term δ is equal to zero, so we neglect all effects of the organic reaction. Let us return our esophageal tissue. Gauss’s theorem(Feynman, Leighton et al. 1963) tell us that the total resultant diffusion flux from an iodine-stain area is equal to the sum of the fluxes out of each cell(cube) because each cubic is an infinitesimal fluid element(Figure E), so we form the sum:

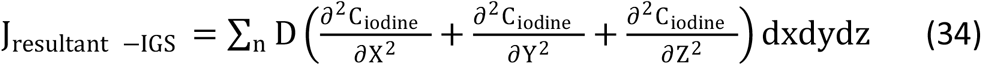

Where n is the number of cubes.

The diffusion is a continuous process, so:

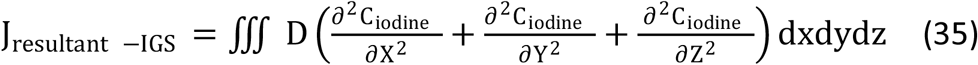

For some researchers’ view(Peng, Li et al. 2020), High-frequency mutation genes encoding epigenetic modification suggest liquid-liquid phase separation may be present in the occurrence or progression of esophageal cancer. Liquid-liquid phase separation can convert into a solid-like state in the pathological state(Dogra, Joshi et al. 2019).it is possible that Iodine cannot diffuse into these tumor cells because their cytoplasm tends to a solid phase. It is not easy to conclude without enough evidence, so we need to prove it with the help of some mathematical logic. Let us first assume that the tumor cell’s cytoplasm tends to the solid phase.

According to Lennard-Jones potential(Israelachvili 2011), the attractive force is dominated in the solid phase, making it difficult to break these molecules’ bonds. When Iodine tried to diffuse into the tumor cell, the concentration gradient will disappear in Y-axis, so we find:

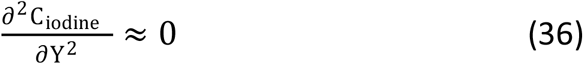

Suppose that we imagine for a moment that Iodine diffuse into the standard epithelial layer above BLSCC(Figure E).when these iodine molecules touch these tumor cells, we have its resultant diffusion flux :

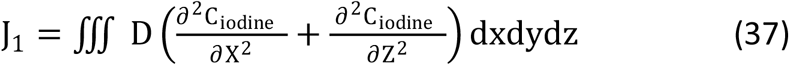

where J_1_ is the resultant diffusion flux of Iodine which diffuses into the standard epithelial layer above BLSCC.

Meanwhile, some iodine molecules diffuse into the standard epithelial layer without BLSCC, so we have its resultant diffusion flux:

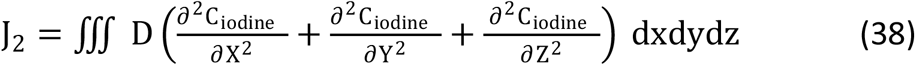

where J_2_ the resultant diffusion flux of Iodine which diffuses into the standard epithelial layer without BLSCC.

Eq(36) means:

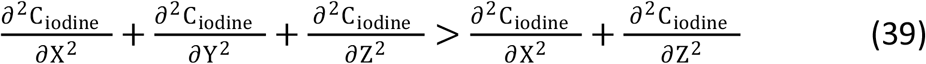

According to the domination property of triple integrals(Thomas, Weir et al. 2008), we have:

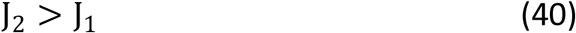

Use Eq(32) and the definition of average value a function in space, we have:

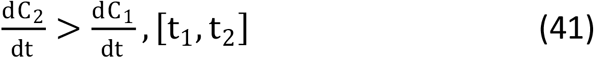

where C_1_ is the concentration of Iodine, which diffuses into the normal epithelial layer above BLSCC; C_2_ is the Iodine concentration, which diffuses into the normal epithelial layer without BLSCC. [t_1_, t_2_] is the diffusion of Iodine for a certain period.

Similar, definite integrals also have domination property, so:

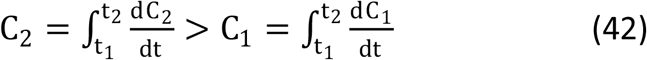

Eq(15) tell us C_iodine −s_ is positively associated with C_omplex −s_, so Eq(42) means:

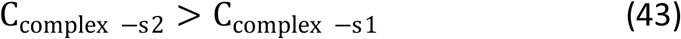

where C_omplex −s2_ is the concentration of Iodine- glycogen complex without BLSCC; C_omplex −s1_ is the Iodine concentration-glycogen Complex above BLSCC.

Let C_omplex −s_ approach M, so C_iodine −s_ tend to +∞ base on Eq(15). Usually, Lugol’s solution needs to spray enough Iodine through a spray catheter, compare with the large value of C_iodine −s_. C_omplex −s_ is in low concentration so that we can use Beer-Lambert law safely in our model.

According to Eq(1), it is logical that the efficacy of Lugol’s staining is both determined by l_esophagus_ and C_complex_. However, the second law of thermodynamics tells us Iodine tends to the highest entropy (uncertainty is largest), so Iodine will have the same concentration at the end. In short, it seems l_esophagus_ only dominates the efficacy of Lugol’s staining. If C_omplex −s_ do not approach M, we also come to the same conclusion.

Can we answer our questions in the current steps? The answer is no. Squamous dysplasia is usually accompanied by dilation and elongation of the interpupillary capillary loop(IPCL)(Kumagai, Toi et al. 2010, Oyama, Inoue et al. 2017). That brings us to another variable we need to consider: the fluid dynamics characteristics of Iodine in IPCL. Compare with the iodine diffusion of cells, and we need to add convective fluxes for IPCL.

Letting C _iodine_ μ be the convective fluxes at an infinitesimal fluid element’s top edge along the x-axis in IPCL and 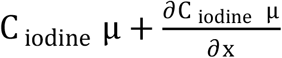 be the convective fluxes at an infinitesimal fluid element’s bottom edge along x-axis base on Taylor polynomial, according to mass conservation, we write:

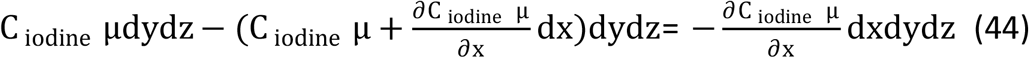

where 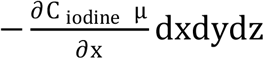 is the net convective fluxes of Iodine at an infinitesimal fluid element along the x-axis for IPCL. Similarly, for the y-axis and z-axis, we write:

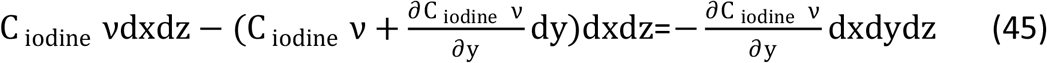

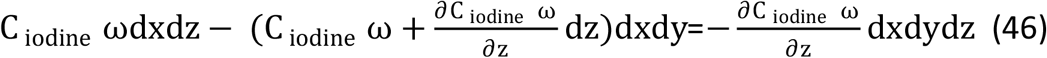

where C _iodine_ ν and C _iodine_ ω are both the convective fluxes at an infinitesimal fluid element’s right and left edge along y-axis and z-axis in IPCL

For an infinitesimal fluid element of IPCL, the concentration variation rate of Iodine is equal to the sum of convection fluxes and diffusion fluxes, therefore:

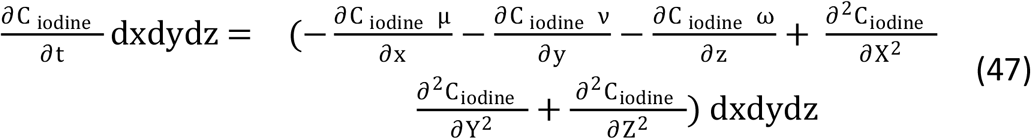

From Eq(33), we know at once:

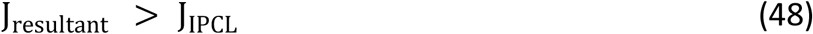

where J_IPCL_ is the concentration variation rate of Iodine at an infinitesimal fluid element of IPCL.

For the same volume, we use the domination property of triple integrals again; we have:

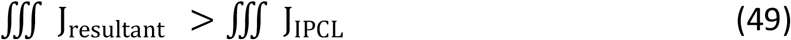

According to the definition of average value a function in space(Thomas, Weir et al. 2008), we get:

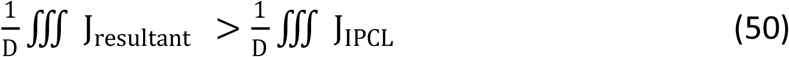

where D is the same volume of iodine diffusion area in both cells (cubic) and IPCL.

Therefore:

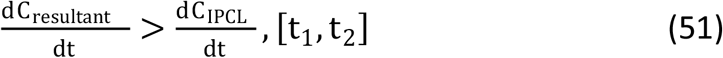

where C_resultant_ is the concentration of iodine in cells; CIPCL is the concentration of iodine in ICPL.

According to the domination property of definite integrals:

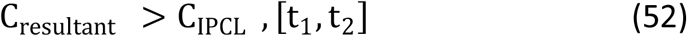

Owing to convective fluxes always exist in a blood vessel, it is clear that” C_resultant_ > C_IPCL_” does not disappear until all iodine molecules are absorbed into the bloodstream. Unlike the iodine diffusion among cells, the diffusion from cells to IPCL always exists during the operational process of Lugol’s Chromoendoscopy. In short, our two questions can be answered by two factors: the thickness of the squamous epithelial cell layer, which contains glycogen granules, and the number of IPCL. In other words, if BLSCC has enough glycogen granules and fewer IPCL, Lugol’s Chromendoscopy does not work. The two factors are consistent with endoscopic characteristics (Zhuang, Chen et al. 2019)and histology structure in BLSCC.

Suppose that the tumor cell’s cytoplasm always keeps liquid phase. According to the clinical observation data show that BLSCC is a common accompaniment to typical squamous cell carcinoma (SCC)(Zhuang, Chen et al. 2019), so, logically, both of them have the same physical characteristics in the hypothesis, and Iodine can diffuse into these cells’ cytoplasm.

Now let us consider the “pink color sign” in squamous cell dysplasia. According to Ishihara et al.’s theory(Ishihara, Kanzaki et al. 2013), “pink color sign” is iodine leak into the esophageal lumen first because of lack of the keratin layer. However, under such assumption, a concentration gradient between the area that is the pink color sign(high-grade dysplasia involves the whole epithelium layer) and its peripheral area (dysplasia do not involve the whole epithelium layer) base on the second law of thermodynamics. When the iodine molecules from the area that are a pink color sign leak into the esophageal lumen first, the iodine molecules of the peripheral area continue to spread the area that is a pink color sign because of the driving force from the concentration iodine gradient. That can lead to two situations: 1. The “pink color sign” is a looping structure; its middle always keeps yellow color. 2. It is impossible to keep a static demarcation line between the area that is a pink color sign and its peripheral area during the whole diffusion process. It is a dynamic changing process. The deduction versus its actual endoscopic characteristics provides a conflicting result. Consequently, the assumption is false. If iodine leaks into IPCL first, we come to the same conclusion.

Let us come back to our first hypothesis: the tumor cell’s cytoplasm tends to a solid phase. Suppose that the number of iodine molecules maintains constant in Iodine unstained and stained lesions. For infinitesimal fluid element on the surface of mucosa with the pink color sign, we have:

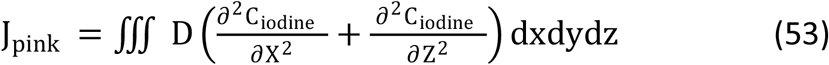

where J_pink_ is the resultant diffusion flux of Iodine on the surface of mucosa with the pink color sign.

For infinitesimal fluid element on the surface of mucosa without the pink color sign, the assumption above tell us the diffusion flux has not existed at the infinitesimal fluid element top edge along the y-axis, so we have

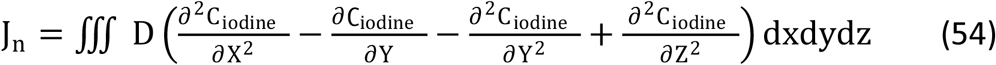

where J_n_ the resultant diffusion flux of Iodine on the surface of mucosa without the pink color sign.

Obviously

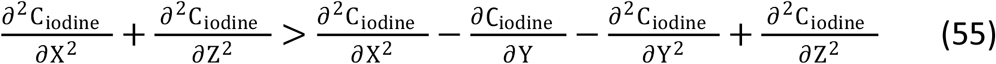

Therefore:

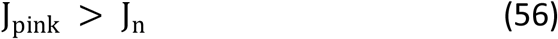

We use the definition of average value a function in space and the domination property of definite integrals again, we have:

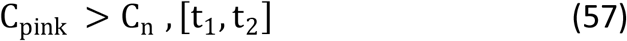

where C_pink_ is the concentration of Iodine on the surface of mucosa with the pink color sign; C_n_ is the concentration of Iodine on the surface of mucosa without the pink color sign.

According to the second law of thermodynamics, it is clear that Iodine moves from the surface of mucosa with a “pink color sign” to the surface of mucosa without the pink color sign. Further, the” 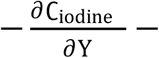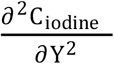” term of Eq(54) tells us iodine molecules continue to leak into the epithelial layer. High-grade intraepithelial neoplasia is usually accompanied by expansion, distortion pipe diameter of IPCL(Kumagai, Toi et al. 2010), so the convection fluxes can accelerate the process base on Eq(52). From the point of view of histology, IPCL does not interfere with the diffusion of Iodine on the mucosal surface because it cannot extend to the top mucosal surface. In conclusion, the normal esophagus epithelium layer continues to absorb iodine molecules on the mucosal surface, leading to a shallow concentration of Iodine on the mucosal surface. When the upper half of epithelium is replaced by neoplastic cells in a certain area, a “pink color sign” can happen because only a few iodine molecules remained in its area. Our model hypothesizes that” the tumor cell’s cytoplasm tends to solid phase” is true.

Suppose that number of iodine molecules is not less than the transport protein on the cell membrane. It is clear that J_top edge −x_ is invariable, and the concentration of iodine molecules is saturation on the top edge of the cell membrane; however, Eq(3) and Eq(4) tell us:

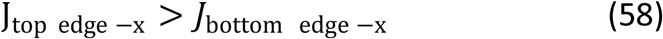

It suggests the number of iodine molecules is less than the transport protein on the bottom edge of the cell membrane. Hence, the derivation process is the same as the previously assumed condition after Iodine diffuses into the mucosal layer.

## Discussion

From the fractal dimension point of view(Grizzi, Russo et al. 2005), the tumor is an actual three-dimensional structure, and tumor-associated angiogenesis is on the non-integer dimensions (it is between 2D and 3D). Non-integer dimensions mean the rate of tumor angiogenesis is less than the tumor growth rate. As the tumor overgrows, the energy available per cell will be less and less. Although the Warburg effect (Basan, Hui et al. 2015)may be a solution from the angle of cost-benefit analysis, reprogramming energy metabolism is not enough to solve the conflict between the overgrowth of cancer cells and energy requirement. This conflict inevitably leads to a substantial morphological change of blood vasculature: tortuous, irregular branched and circumferential enlarged et al(Farnsworth, Lackmann et al. 2014), so It is not strange that the degree of micro-vascular irregularity is associated with the invasion depth of superficial squamous cell carcinoma(Oyama, Inoue et al. 2017). the nutrients and oxygen first move into the extra-vascular tissue by the flow of blood(convection) and then diffuse into the tumor tissue with the help of a concentration gradient(Baish, Gazit et al. 1996). For- dimensional diffusion in a plane, the maximum distance of oxygen diffusion is 100um(Secomb, Hsu et al. 1993, Baish, Gazit et al. 1996)(note: the actual thickness of epithelium is at least greater than 200um)(Vieth, Mastracci et al. 2016). Warburg effect means glucose is the principal metabolic fuel of cancer cells, so the maximum distance of glucose diffusion is significant for the energy requirement of cancer cells.

Compare with oxygen’s simple diffusion, Eq(58) tells us the maximum distance of glucose’s facilitated diffusion is minor than 100um because the number of membrane proteins limits its concentration gradient. Come back to the histological structure of Basal layer-type squamous cell carcinoma in situ; the lamina propria usually contains many vessels (Mills 2012). The basement membrane has many pore(Lv, Chen et al. 2014), so these histological characteristics mean glucose can diffuse into the epithelium through the basement membrane without the help of the interpupillary capillary loop(note: the maximum distance of glucose is facilitated diffusion may be further lessened by porous medium). When the distance from the basement membrane to the cancer cell is less than the maximum distance of glucose diffusion, it is logical that tumor-associated angiogenesis is unnecessary at that distance. For BLSCC, the histological characteristics suggest that many parts of it are located within the maximum distance of glucose diffusion. These components contain a fewer number of interpupillary capillary loops. However, it is essential to note that tumor-associated angiogenesis depends on the maximum distance of glucose diffusion and the glycolysis rate of tumor cell(Chen, Yi et al. 2021). A histological perspective means tumor-associated angiogenesis also depends on the degree of cytological atypia(Figure D). In brief, the essential prerequisite for tumor-associated angiogenesis is:

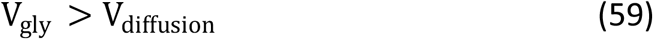

where V_gly_ is the glycolysis rate of tumor cells, V_diffusion_ is the glucose rate that diffuses into the tumor cell from lamina propria.

According to Eq(1) and Eq(2), the distance of glucose diffusion and the concentration of glucose present negative relativity, so Eq(59) tells us tumor-associated angiogenesis is the integration effect between the distance of glucose diffusion and the glycolysis rate of tumor cell. When the glycolysis rate is faster, the degree of cytological atypia is more prominent, and tumor cell without IPCL is closer to the basement membrane. Metabolomics study(Chen, Yi et al. 2021) suggests the increased glycolysis rate of tumor cells is a gradual process, so it means the growth of tumor cells without IPCL exists in the specific time frame. Compared with the evolution of typical SCC, BLSCC has fewer IPCL in the specific time frame.

Moreover, These BLSCC components usually locate more close to the basement membrane to replace enough cells containing glycogen. To detect BLSCC, the sensitivity of Lugol’s staining may be finally determined by the maximum distance of glucose diffusion and the degree of cytological atypia. When IPCLs are few on this condition, some components of BLSCC also may be able to escape the detection of Narrow-band imaging with magnifying endoscopy.

Similarly, low-grade squamous intraepithelial neoplasia/dysplasia (LGIN) (Board 2019) maybe not also be detected by Lugol’s chromoendoscopy and Narrow-band imaging with magnifying endoscopy on the same condition. Compare with BLSCC on the condition, LGIN may not have a vast impact on the prognosis of squamous cell carcinoma over some time because most of it lacks early infiltration (Chen, Xu et al. 2017)from epithelium to submucosa. This difference may stem from the difference in reprogramming energy metabolism. Metabolomics study (Chen, Yi et al. 2021)also showed glycolysis rate of high grade squamous intraepithelial neoplasia might be higher than low grade in cervical squamous intraepithelial lesion whose structure is similar to the esophagus. It suggests microenvironment of BLSCC may be more acidic than LOGIN in the esophagus (diffusion of hydrogen ion). Low PH condition is usually associated with high expression of matrix metalloproteases and invasive tumor front(Rohani, Hao et al. 2019). Owing to BLSCC close to basement membrane on the condition, it is logical that BLSCC tends to infiltration at an earlier stage with completely missing basement membrane(Chen, Xu et al. 2017). besides, when BLSCC and LGIN above the maximum distance of nutrients, the degree of irregularity in the BLSCC’s micro-vascular morphology is more severe than LOGIN from the energy requirement point of view.

According to the classification of intermolecular forces’ different physical or chemical origin(Israelachvili 2011), it is clear that reprogramming energy metabolism can affect intermolecular forces inside the cell are purely electrostatic origin. Research(Alshareedah, Kaur et al. 2019) shows that the electrostatic force plays a crucial role in driving the formation of Liquid-liquid phase separation. It is possible that reprogramming energy metabolism can break phase separation of enzymes’ homeostasis with the help of low PH condition(Prouteau and Loewith 2018, Dignon, Best et al. 2020), so the Warburg effect suggests BLSCC’s cytoplasm may be closer to solid-phase than LGIN. Come back to our model, it suggests iodine molecules may permeate cells of LGIN more easily than BLSCC because of their different degree of Warburg effect. The inference will cause a decline of the term” C_iodine −s_” so that “C_omplex −s_“ is reduced base on Eq(15). According to Eq(1), LGIN is easier to undergo color fading than BLSCC in the same condition of Lugol’s staining. From a data science perspective for detecting early cancer of the esophagus, LOGIN may have a higher positive rate than BLSCC for Lugol’s Chromoendoscopy. Clinical data also hints at this trend(Dawsey, Fleischer et al. 1998). on the other side of the spectrum, Lennard-Jones potential suggests cytoplasm’s solid phase could prevent oxygen’s simple diffusion, so it means cancer cells have to choose anaerobic metabolism. It is more than likely that reprogramming energy metabolism and LLPS are mutually reinforcing relationships.

Although BLSCC is regarded as a kind of high-grade squamous dysplasia(Board 2019), a question remains: do BLSCC have an independent evolution process relative to typical superficial esophageal carcinoma? The cytological resemblance between them is remarkable. Most cases of BLSCC usually coexist with typical SCC components or LGIN components(Zhuang, Chen et al. 2019). These histological characteristics suggest BLSCC and typical superficial esophageal carcinoma might share the same evolution way even though the lamina propria mucosae is easier to be infiltrated by BLSCC than LGIN. Still, in the absence of any further experimental evidence, we cannot answer this question fully. From a data science point of view, it is a potential choice to construct a phylogeny tree(Schwartz 2008) base on data from next-generation sequencing(NGS) and an algorithm for answering the question. Phylogeny tree is involved in depicting the evolutionary relationship among species and exploring various tumor lineage(Coorens, Farndon et al. 2020). Logically, we can construct a phylogeny tree among BLSCC, LGIN, typical superficial esophageal carcinoma, and regular esophageal tissue base on specific algorithms and data from NGS. However, inter-observer variation in the pathological diagnosis of BLSCC and LGIN (Zhuang, Chen et al. 2019) may bring difficulty to phylogeny analysis, so how to raise pathological diagnostic accurate is the key to construct the phylogeny tree.

## Conclusion

In conclusion, through an analysis of diffusion dynamics of Lugol’s chromoendoscopy in Basal layer-type squamous cell carcinoma in situ, our model reveals that the efficacy of Lugol’s chromoendoscopy is determined by the thickness of the squamous epithelial cell layer, which contains glycogen granules and the number of IPCL. When some components of BLSCC near the basement membrane, Lugol’s chromoendoscopy’s effect should weaken because their energy requirement does not only rely on tumor angiogenesis, and their unique histological structure does not allow them to break squamous epithelial cell layer which contains glycogen granules. By extrapolating iodine diffusion from the “pink color sign,” it is possible that the cytoplasm of BLSCC and typical superficial esophageal carcinoma tend to solid-phase if they both share the same evolution path.

## Reference

Agresti, A. and C. Franklin (2018). Statistics the art and science of learning from data, Pearson Education Limited.

Alberti, S., et al. (2019). “Considerations and challenges in studying liquid-liquid phase separation and biomolecular condensates.” Cell 176(3): 419–434.

Alshareedah, I., et al. (2019). “Interplay between short-range attraction and long-range repulsion controls reentrant liquid condensation of ribonucleoprotein–RNA complexes.” Journal of the American Chemical Society 141(37): 14593–14602.

Anderson, J. D. and J. Wendt (1995). Computational fluid dynamics, Springer.

Baish, J. W., et al. (1996). “Role of tumor vascular architecture in nutrient and drug delivery: an invasion percolation-based network model.” Microvascular research 51(3): 327–346.

Basan, M., et al. (2015). “Overflow metabolism in Escherichia coli results from efficient proteome allocation.” Nature 528(7580): 99–104.

Board, W. C. T. E. (2019). Digestive System Tumours, International Agency for Research on Cancer.

Chen, G., et al. (2017). “The expression of podoplanin protein is a diagnostic marker to distinguish the early infiltration of esophageal squamous cell carcinoma.” Oncotarget 8(12): 19013.

Chen, X., et al. (2021). “Metabolomics study reveals the potential evidence of metabolic reprogramming towards the Warburg effect in precancerous lesions.” Journal of Cancer 12(5): 1563.

Cho, J. W., et al. (2014). “Lymph node metastases in esophageal carcinoma: an endoscopist’s view.” Clinical endoscopy 47(6): 523.

Coorens, T. H., et al. (2020). “Lineage-Independent Tumors in Bilateral Neuroblastoma.” New England Journal of Medicine 383(19): 1860–1865.

Dawsey, S. M., et al. (1998). “Mucosal iodine staining improves endoscopic visualization of squamous dysplasia and squamous cell carcinoma of the esophagus in Linxian, China.” Cancer: Interdisciplinary International Journal of the American Cancer Society 83(2): 220–231.

Dignon, G. L., et al. (2020). “Biomolecular phase separation: From molecular driving forces to macroscopic properties.” Annual review of physical chemistry 71: 53–75.

Dogra, P., et al. (2019). “Intermolecular Charge-Transfer Modulates Liquid–Liquid Phase Separation and Liquid-to-Solid Maturation of an Intrinsically Disordered pH-Responsive Domain.” Journal of the American Chemical Society 141(51): 20380–20389.

E Nunez-Anita, R., et al. (2011). “Peroxisome proliferator-activated receptors: role of isoform gamma in the antineoplastic effect of iodine in mammary cancer.” Current cancer drug targets 11(7): 775–786.

Endeward, V. (2012). “The rate of the deoxygenation reaction limits myoglobin-and hemoglobin-facilitated O2 diffusion in cells.” Journal of Applied Physiology 112(9): 1466–1473.

Farnsworth, R. H., et al. (2014). “Vascular remodeling in cancer.” Oncogene 33(27): 3496–3505.

Ferner, R. E. and J. K. J. B. j. o. c. p. Aronson (2016). “Cato Guldberg and Peter Waage, the history of the Law of Mass Action, and its relevance to clinical pharmacology.” British journal of clinical pharmacology 81(1): 52–55.

Feynman, R. P., et al. (1963). The Feynman Lectures on Physics, Addison-Wesley Publishing Company.

Freitag, C., et al. (1999). “Esophageal dysplasias are detected by endoscopy with Lugol in patients at risk for squamous cell carcinoma in southern Brazil.” Dis Esophagus 12(3): 191–195.

Funakawa, K., et al. (2015). “Effect of endoscopic submucosal dissection for superficial esophageal neoplasms and risk factors for postoperative stricture.” Medicine 94(1).

Goda, K., et al. (2015). “Narrow-band imaging magnifying endoscopy versus Lugol chromoendoscopy with pink-color sign assessment in the diagnosis of superficial esophageal squamous neoplasms: a randomised noninferiority trial.” Gastroenterology research and practice 2015:639462.

Grizzi, F., et al. (2005). “Quantitative evaluation and modeling of two-dimensional neovascular network complexity: the surface fractal dimension.” BMC cancer 5(1): 1–9.

Hashimoto, C. L., et al. (2005). “Lugol’s dye spray chromoendoscopy establishes early diagnosis of esophageal cancer in patients with primary head and neck cancer.” The American journal of gastroenterology 100(2): 275–282.

Ishihara, R., et al. (2013). “Pink-color sign in esophageal squamous neoplasia, and speculation regarding the underlying mechanism.” World journal of gastroenterology 19(27): 4300.

Ishihara, R., et al. (2010). “Prospective evaluation of narrow-band imaging endoscopy for screening of esophageal squamous mucosal high-grade neoplasia in experienced and less experienced endoscopists.” Diseases of the Esophagus 23(6): 480–486.

Israelachvili, J. N. (2011). Intermolecular and surface forces, Academic press.

Kanamoto, A., et al. (2000). “Clinicopathological study of multiple superficial oesophageal carcinoma.” The British journal of surgery 87(12): 1712–1715.

Kumagai, Y., et al. (2010). “Angiogenesis in superficial esophageal squamous cell carcinoma: magnifying endoscopic observation and molecular analysis.” Digestive Endoscopy 22(4): 259–267.

Lv, J., et al. (2014). “Promoting epithelium regeneration for esophageal tissue engineering through basement membrane reconstitution.” ACS applied materials interfaces 6(7): 4954–4964.

Manner, H., et al. (2013). “Efficacy, safety, and long-term results of endoscopic treatment for early stage adenocarcinoma of the esophagus with low-risk sm1 invasion.” Clinical gastroenterology and hepatology : the official clinical practice journal of the American Gastroenterological Association 11(6): 630–635.

Mayerhöfer, T. G., et al. (2020). “The Bouguer - Beer - Lambert law: Shining light on the obscure.” Chemphyschem : a European journal of chemical physics and physical chemistry 21(18): 2029.

Mills, S. E. (2012). Histology for Pathologists, Wolters Kluwer Health.

Moulay, S. (2013). “Molecular iodine/polymer complexes %J Journal of Polymer Engineering.” Journal of Polymer Engineering 33(5): 389–443.

Mourão, M. A., et al. (2014). “Connecting the dots: the effects of macromolecular crowding on cell physiology.” Biophysical journal 107(12): 2761–2766.

Oyama, T., et al. (2017). “Prediction of the invasion depth of superficial squamous cell carcinoma based on microvessel morphology: magnifying endoscopic classification of the Japan Esophageal Society.” Esophagus 14(2): 105–112.

Peng, L., et al. (2020). “From start to end: phase separation and transcriptional regulation.” J Biochimica et Biophysica Acta -Gene Regulatory Mechanisms: 194641.

Prouteau, M. and R. Loewith (2018). “Regulation of cellular metabolism through phase separation of enzymes.” Biomolecules 8(4): 160.

Rohani, N., et al. (2019). “Acidification of tumor at stromal boundaries drives transcriptome alterations associated with aggressive phenotypes.” Cancer research 79(8): 1952–1966.

Rosen, K. (2011). Discrete Mathematics and Its Applications, McGraw-Hill Education.

Sarbia, M., et al. (2015). “High-grade dysplasia, restricted to the basal cell layer involving the entire esophagus.” World journal of gastroenterology 21(5): 1663.

Schwartz, R. (2008). Biological Modeling and Simulation: A Survey of Practical Models, Algorithms, and Numerical Methods, MIT Press.

Secomb, T. W., et al. (1993). “Analysis of oxygen transport to tumor tissue by microvascular networks.” International Journal of Radiation Oncology* Biology* Physics 25(3): 481–489.

Thomas, G. B., et al. (2008). Thomas’ Calculus, Pearson Addison Wesley.

Van Meer, G., et al. (2008). “Membrane lipids: where they are and how they behave.” Nature reviews. Molecular cell biology 9(2): 112–124.

Vieth, M., et al. (2016). “Epithelial thickness is a marker of gastroesophageal reflux disease.” Clinical Gastroenterology and hepatology 14(11): 1544–1551. e1541.

Wang, G.-Q., et al. (2004). “Long-term results of operation for 420 patients with early squamous cell esophageal carcinoma discovered by screening.” The Annals of thoracic surgery 77(5): 1740–1744.

Zhuang, X., et al. (2019). “Clinicopathological features and outcomes of esophageal lesions containing a basal layer type squamous cell carcinoma component.” Cancer management and research 11: 8507.

